# Protein kinase Sut1 from the Crenarchaeon *Sulfolobus acidocaldarius* displays tyrosine phosphorylation activity

**DOI:** 10.1101/2024.01.14.552423

**Authors:** Hassan R. Maklad, Sébastien Pyr dit Ruys, Indra Bervoets, Didier Vertommen, Gustavo J. Gutierrez, Wim F. Vranken, Bettina Siebers, Eveline Peeters

## Abstract

Protein phosphorylation is a key cellular signaling mechanism that exists in all life forms. Unlike Bacteria and Eukarya, in which protein phosphorylation has thoroughly been studied, post-translational modification by means of phosphorylation have only been limitedly explored in Archaea. A previous study of the phosphoproteome of the model Crenarchaeon *Sulfolobus acidocaldarius* revealed a widespread occurrence of protein phosphorylation, especially on tyrosine residues. Moreover, several (putative) transcription factors, including AbfR1 and FadR, were previously shown to be phosphorylated on tyrosine residues, a phenomenon that is directly linked to a phosphorylation-mediated regulation of these transcription factors. Despite this potentially important role for tyrosine phosphorylation in *S. acidocaldarius*, to our knowledge, a kinase capable of directly phosphorylating tyrosine residues has not yet been identified in this organism, neither in Archaea as a whole. Here, we identify and characterize a protein kinase in *S. acidocaldarius*, Sut1, which displays tyrosine phosphorylation activity *in vitro* and represents a novel protein kinase family that is widespread in different archaeal organisms.

## Introduction

Reversible protein phosphorylation is a signal transduction mechanism that is exploited by all types of living cells. By modulating the functions of a myriad of proteins involved in virtually all cellular processes, protein phosphorylation enables cells to respond to various types of stimuli for the sake of their fitness and survival [1]. Although extensively studied in eukaryotes and bacteria, little is still known about this process in archaea [2]. Being classified as the “third domain of life” that encompasses microorganisms with diverse and specialized physiologies and metabolic properties (thermophiles, halophiles, and methanogens), a proper understanding of their cellular functioning would benefit from studying protein phosphorylation.

In the model archaeal species *Sulfolobus acidocaldarius*, belonging to the phylum Crenarchaeota, about 37% of the total protein content was found phosphorylated *in vivo*, with 36% of phosphorylation events occurring on tyrosine residues [3]. This underscores the importance of tyrosine phosphorylation in this organism. However, no tyrosine kinase had thus far been identified in archaea, which evokes questions about the source of these tyrosine phosphorylation events.

Like other archaea, *Sulfolobales* harbor many homologs of eukaryotic-type protein kinases (ePKs) [2]. More specifically, the genome of *S. acidocaldarius* encodes at least 10 putative protein kinases, most of which are classified as either typical or atypical Hanks-type kinases. Typical kinases are characterized by the presence of 12 conserved subdomains that harbor specific conserved residues, which play critical functional roles in their catalytic reaction. Among these subdomains are a nucleotide-binding loop, typically with the sequence “GXGXXG”, which mediates the interaction with ATP (subdomain I) and a catalytic loop that contains an invariant catalytic aspartate residue directly involved in phosphoryl transfer (subdomain VIb). Additionally, there is a metal-binding loop or “DFG” loop (subdomain VII) with a conserved aspartate required for anchoring metal cations, typically Mg^+2^. Canonical Hanks-type protein kinases contain an activation loop that is located between the “DFG” motif and the highly conserved “APE” motif (subdomain VIII). In these kinases, phosphorylation occurring on the activation loop results in modulation of kinase activity [4,5]. Most of the kinases in *S. acidocaldarius* are predicted or shown to have serine and threonine phosphorylation activity [2].

In contrast to archaea, protein kinases targeting tyrosine residues were previously identified in bacteria [6-8]. Bacterial tyrosine kinases are characterized by the presence of conserved Walker motifs, namely a Walker A motif “GXXXXGKT/S”, Walker A’ motif “DAD” and Walker B motif “DTPP” [6], in addition to a tyrosine-rich cluster at the C-terminus [7-8].

Here, we report the identification of a novel type of protein kinases, represented by a prototypical kinase from *S. acidocaldarius*, named Sut1. We present an initial characterization of Sut1 kinase activity *in vitro* using kinase assays, western blotting and mass spectrometry. Finally, we implement a phosphoproteomic analysis, aimed to determine Sut1 substrate(s) *in vivo*.

## Results

### Sut1 from S. acidocaldarius represents a novel kinase family in archaea

One of the putative protein kinases encoded in the genome of *S. acidocaldarius*, encoded by gene *Saci_1289*, does not show the typical Hanks-type motifs, although it was annotated as a Ser/Thr protein kinase (**Supplementary Figure S1**). In addition, it lacks the signature of catalytic subdomain VIb, typified by the motif “HRDxxxN”, which is invariably conserved in other Hanks-type kinases. Thus, the *Saci_1289*-encoded protein does not belong to the Hanks-type protein kinase family.

We screened other protein kinases for potential homology, including selected candidates from the bacterial tyrosine kinase family BY, namely CpsD from *Streptococcus pneumoniae*, CapB from *Staphylococcus aureus* and PtkA from *Mycobacterium tuberculosis* (**Supplementary Figure S2**), revealing that the *Saci_1289*-encoded protein harbors motifs corresponding to typical BY motifs, specifically putative Walker A “GKT” and Walker A’ “DxD” motifs. Additionally, the protein encoded by *Saci_1289* harbors a C-terminal tyrosine-rich cluster comprised of 3 tyrosine residues in a 10-amino acid stretch, resembling to some extent the pattern of tyrosine-rich clusters in bacterial BY kinases [8]. In contrast, the highly conserved catalytic Walker B motif is missing in Saci1289 (**Supplementary Figure S2**). While these observations do not predict to unambiguously classify this archaeal protein as a BY-family tyrosine kinase, they suggest that it is a protein kinase of a novel family that might have a tyrosine phosphorylation activity. Based on this assumption, and on the confirmation of its activity as described below, we propose to name the protein encoded by *Saci_1289* as *<u>Su</u>lfolobus* <u>t</u>yrosine kinase 1, Sut1.

To investigate the conservation of Sut1 in Archaea, a BLAST analysis was conducted (**Figure 1, Supplementary Figure S3**). This revealed that Sut1 is not exclusive to *Sulfolobus acidocaldarius* and that homologs are present in other Crenarchaeota and other unrelated archaeal phyla, such as Thaumarchaeota. For example, a Sut1 homolog with 34% sequence identity is present in the mesophilic ammonia-oxidizer *Nitrosopumilus* spp, which has a very different lifestyle with respect to the thermoacidophilic *S. acidocaldarius*. Upon comparing sequences of different archaeal Sut1 homologs, conserved motifs are noticeable (**Figure 1, Supplementary Figure S3**), and with respect to other characterized protein kinase families, putative catalytic motifs were predicted (**Supplementary Figure 2**).

**Figure 1.**
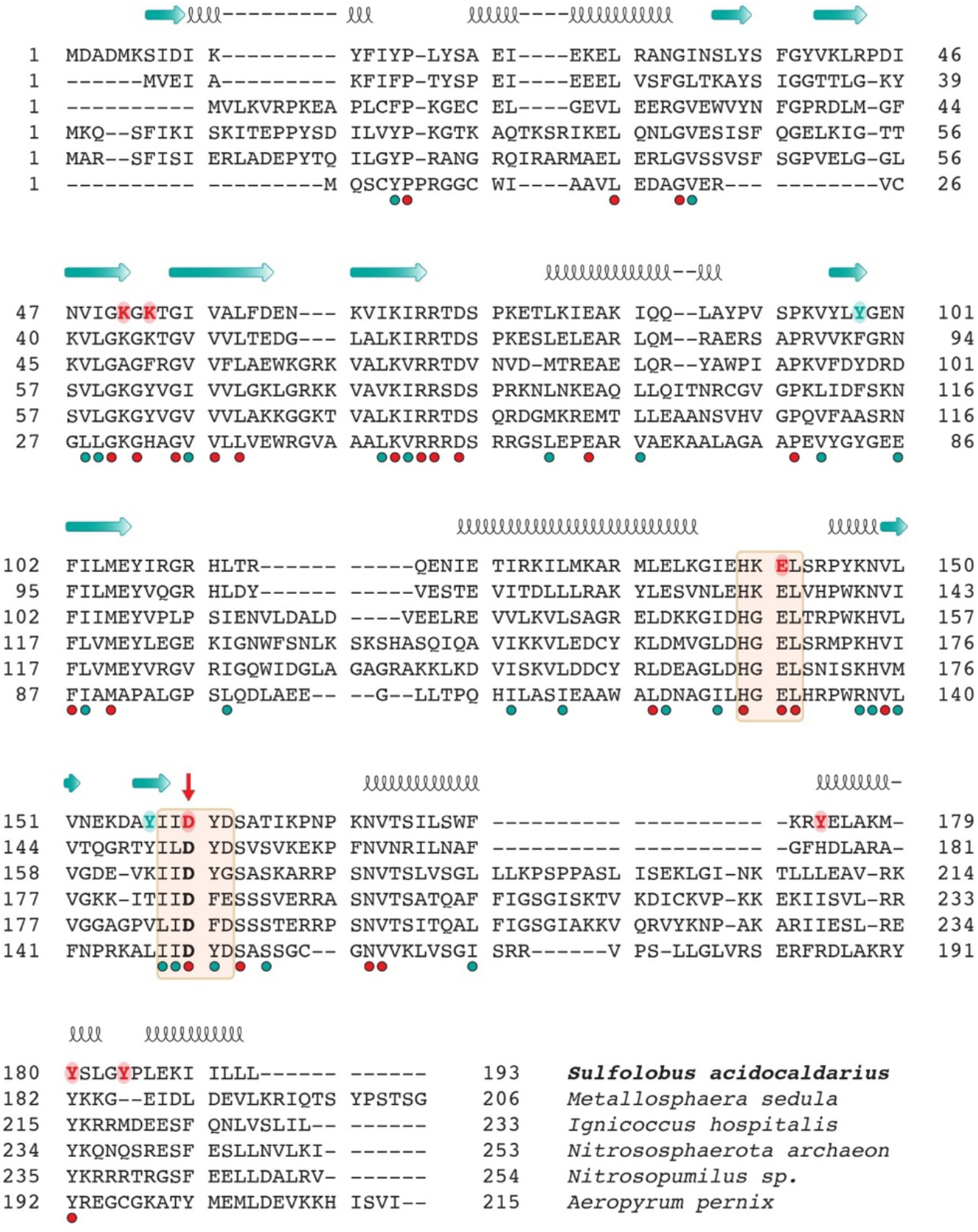
Protein sequence alignment of Sut1 from *S. acidocaldarius* with selected homologs from other archaea *Metallosphaera sedula, Ignicoccus hospitalis*, an uncultured Nitrososphaerota archaeon from the Thaumarchaota phylum, *Nitrospumilus* sp. and *Aeropyrum pernix*. Sut1 residues selected for mutagenesis analysis are depicted in red. Identical and similar residues are annotated with green and red dots, respectively. The red arrow points to the highly conserved catalytic residue Asp160. Predicted secondary structure elements are annotated with arrows for β-strands and spiral helices for α-helices.

### Structural model of Sut1

To gain insights into the structure of Sut1 and its homologs, a threedimensional structural model was retrieved from AlphaFold [9] (**Figure 2**). The structural model exhibits a typical protein kinase fold, with N and C-lobes and a central cleft in between, where phosphotransfer occurs (**Figure 2A**). The N-lobe is composed of 9 antiparallel β-strands and 2 α-helices, whereas the C-lobe is predominately α-helical.

**Figure 2.**
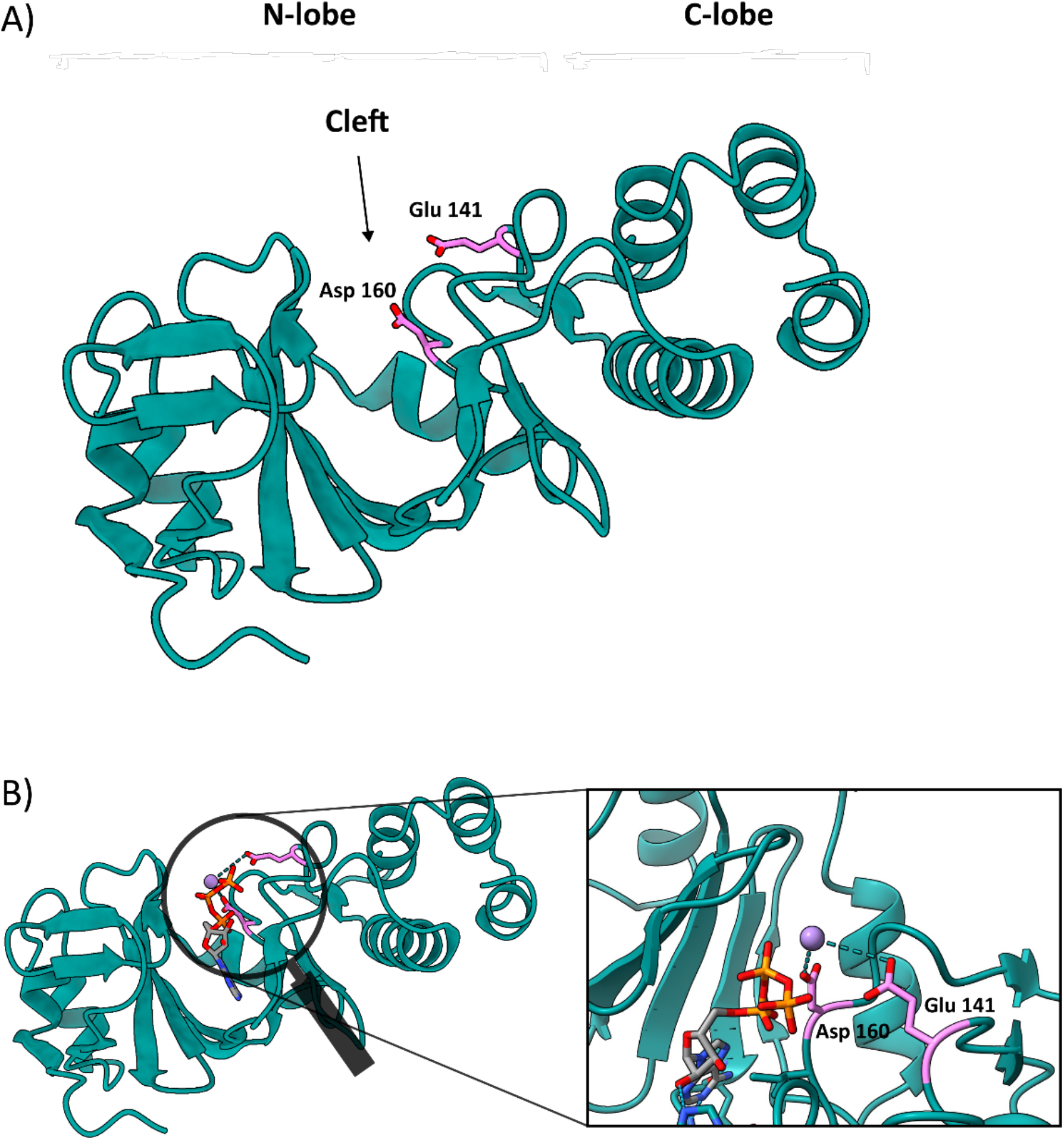
Structural model of Sut1. **A**) Model of Sut1 predicted by AlphaFold (Confidence level: very high (pLDDT > 90)), which shows an N- and C-lobe with a central cleft in between, with the catalytic Glu141 and Asp160 are shown in stick representation in magenta. **B**.The structural model of Sut1 with ligands Mn^2+^ and ATP bound as predicted by Alphafold Filled (AF-Q4J999-F1-model). To the right, a zoom of the catalytic site is shown, highlighting the interactions of Glu141 and Asp160 with Mn^2+^ and ATP.

The position of ligands and cofactors in the structural model was predicted by AlphaFill, which employs sequence and structure similarity to ‘transplant’ ‘missing’ small molecules and ions from protein structural models [10]. Interestingly, a manganese Mn^2+^ cation was predicted to bind in the cleft (RMSD 0.05) through direct interactions with Asp160 and Glu141. Additionally, ATP and ADP molecules were also predicted to be embedded in the cleft (RMSD 1.8) and to establish interactions with Glu141 and juxtaposed residues. The relative positioning of these specific residues in the central cleft suggests their involvement in catalysis (**Figure 2B**).

Besides being highly conserved in archaeal homologs, Asp160 and Glu141 are facing each other in Sut1 (**Figure 2B**). The motif “IIDYD”, to which Asp160 belongs, can be predicted to be a cation-binding motif, thus functionally corresponding to the highly conserved catalytic subdomain VIb with the motif “HRDxxxxN” in Hanks-type kinases [11]. Furthermore, the motif “H^K^/_G_EL”, containing Glu141, can be predicted to correspond to the “DFG” motif in Hanks-type kinases [11], in which it is highly conserved and plays a pivotal role in ATP docking and in coordinating Mg^2+^ binding. In contrast to Hanks-type kinases, the aspartate corresponds to a glutamate in Sut1. The observation that Asp160 and Glu141 are located close to each other in the Sut1 structural model (**Figure 2A** and **B**), is in line with the typical protein kinase fold, in which the DFG and HRD motifs are oppositely positioned in the central cleft [11]. This makes involvement of the aforementioned conserved motifs in ATP binding and catalysis of phosphotransfer likely.

### Sut1 displays kinase activity in vitro

Previously, the Sut1 homolog in *S. islandicus*, encoded by *SiRe_1057* and displaying a sequence identity of 58% with *S. acidocaldarius* Sut1, was tested in an *in vitro* kinase assay but did not show any phosphorylation activity [12]. In this work, we cloned and heterologously expressed the *S. acidocaldarius* Sut1 in *E. coli*, followed by a His-tag affinity purification. Subsequently, given that protein kinases typically perform autophosphorylation, the purified protein was tested for autophosphorylation activity in an *in vitro* phosphorylation assay.

Initial radioactive kinase assays showed very weak phosphorylation signals, suggesting a autophosphorylation activity of *S. acidocaldarius* Sut1 (**Figure 3, Supplementary Figure S4**). We then aimed to optimize the observed *in vitro* activity by exploring different assay conditions. First, we tested two reaction buffer compositions: buffer A (50 mM Tris-HCl [pH 7.8], 10 mM MgCl_2_, 10 mM NaCl and 1mM DTT) [11] and buffer B (75 mM MES [pH 6.5], 45 mM KCl, 3 mM MgCl_2_, 3 mM MnCl_2_) [13] with different kinase concentrations. In buffer A, the autophosphorylation signal was about three times higher relative to buffer B and it directly correlated with the kinase concentration (**Supplementary Figure S4**). Next, different reaction temperatures were assayed (**Figure 3A**). In earlier previous studies, *in vitro* phosphorylation assays of *Sulfolobus* kinases were mostly conducted at 55°C [12,13]. However, this assay demonstrated that Sut1 displays around 20% higher activity at a higher, more physiologically relevant temperature of 70°C. While autophosphorylation activity was highest at 70°C, the protein ultimately degraded when incubated at 80°C (**Figure 3A**).

**Figure 3.**
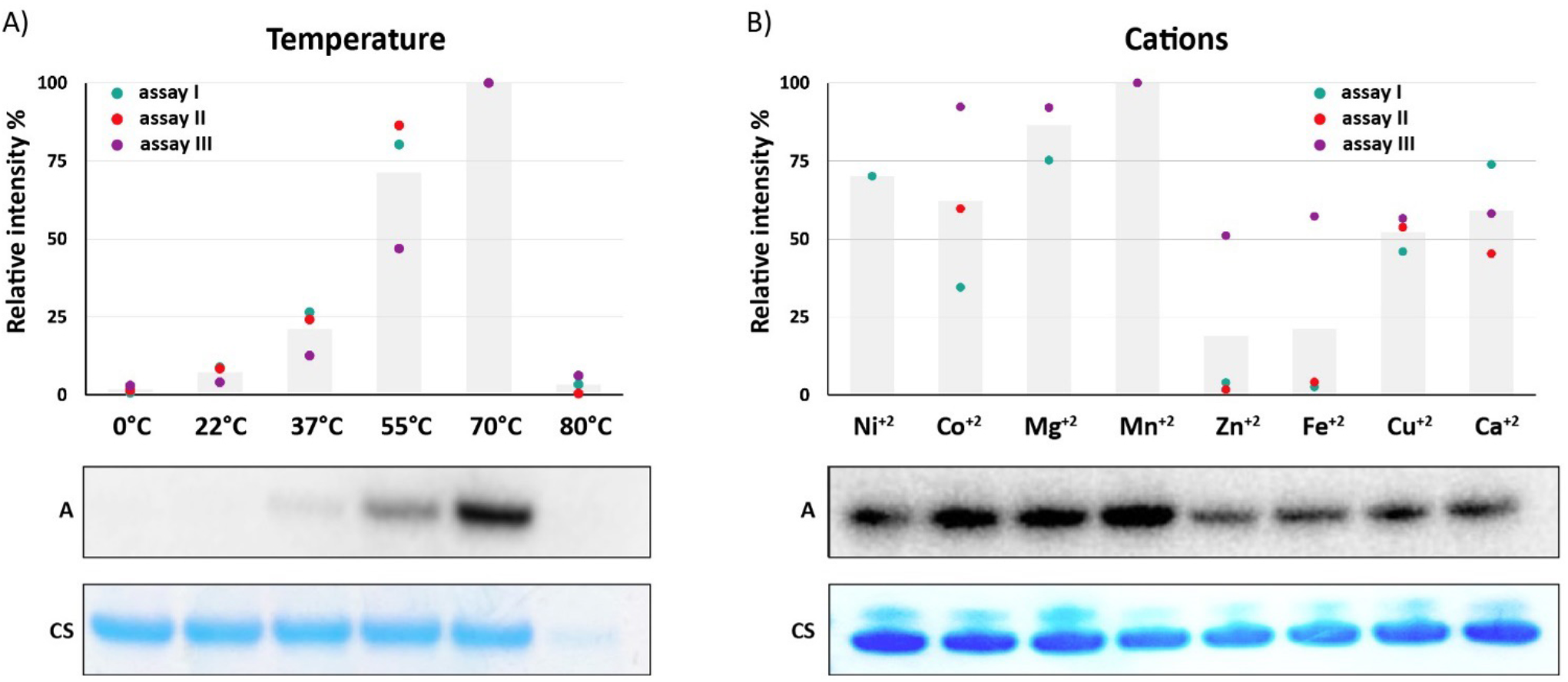
Sut1 displays autophosphorylation activity *in vitro*. **A**) Effect of temperature on Sut1 autophosphorylation activity. A representative Coomassie-stained (CS) gel and autoradiograph (A) are shown. **B**. Effect of different cations on Sut1 autophosphorylation activity. A representative Coomassie-stained (CS) gel and autoradiograph (A) are shown. The histograms represent the average of three independent assays (indicated by coloured dots). Relative intensities were estimated using the Personal Molecular Imager PMI software (BioRad).

Divalent cations are essential cofactors for protein kinases as they play a crucial role in phosphotransfer catalysis. They interact with the negatively charged beta and gamma phosphate groups of ATP, thus stabilizing its binding to the kinase catalytic site [14]. The effect of several cations on Sut1 autophosphorylation activity was therefore tested (**Figure 3B**). We found that Sut1 activity was higher in the presence of Mn^2+^, Co^2+^ or Mg^2+^ relative to the other cations. This observation is consistent with the previously characterized *Sulfolobus* kinases, which also display a preference for Mg^2+^ or Mn^2+^ as cofactors [15,16]. When assaying Sut1 during different incubation times, the activity was maximal after 1 hour of incubation time. The specific activity of Sut1 autophosphorylation was calculated as 166 pmol/mg/min, which is relatively low, suggesting that other relevant cofactors (for example, other proteins that are required for Sut1 activation) are likely missing in the *in vitro* assay. This might also explain the lack of *in vitro* phosphorylation activity detection for Sut1 homologs in previous studies [12].

### Sut1 specifically phosphorylates tyrosine residues

To investigate the hypothesis that Sut1 phosphorylates tyrosine residues, the kinase was challenged with a poly-(Glu_4_-Tyr) peptide substrate, which is a generic tyrosine kinases substrate, in presence of [γ-^32^P] ATP [17,18] (**Figure 4A**). Clear phosphorylation signals were detected for the poly-(Glu_4_-Tyr) substrate, creating a smeared signal in the autoradiograph, with the intensities proportional to the polypeptide concentration. Additionally, autophosphorylation appeared to be stimulated in presence of higher concentrations of the substrate. As a negative control, a similar assay was performed without adding Sut1, for which no phosphorylation signals of the substrate were observable, indicating that the phosphorylation signals are exclusively due to Sut1 phosphorylation activity. Moreover, a similar assay was performed for the well-characterized Ser/Thr kinase from *S. acidocaldarius*, ArnD, which is known to phosphorylate only serine and threonine residues [13]. In this assay, signals were only detected for autophosphorylation but not for the phosphorylation of the poly-(Glu_4_-Tyr) substrate (**Figure 4A**). These observations further strengthen the assumption that Sut1 is capable of phosphorylating tyrosine residues.

**Figure 4.**
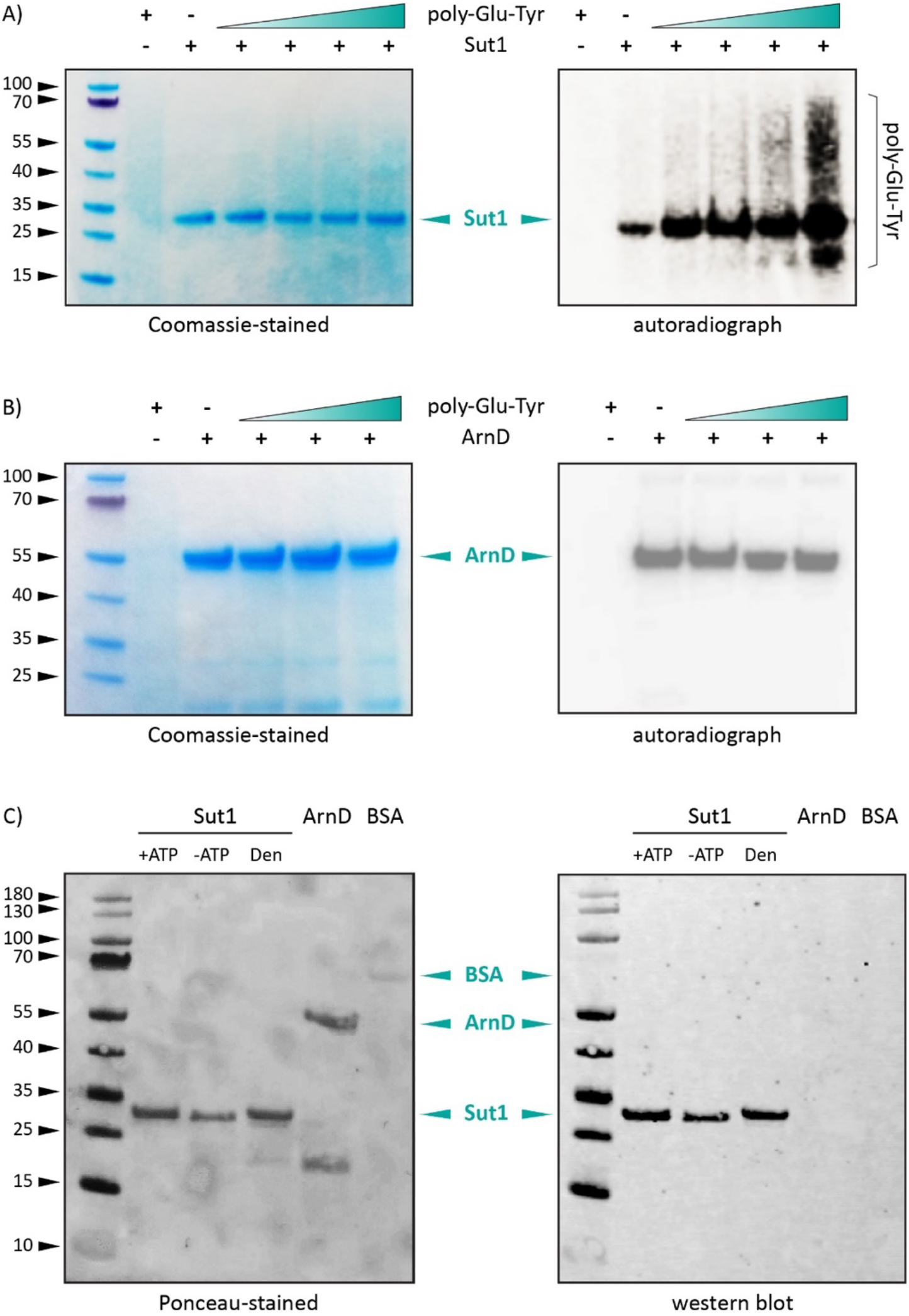
Sut1 phosphorylates tyrosine residues. *In vitro* phosphorylation assay of **A**) Sut1 (23.6 kDa) and **B)** ArnD (47 kDa) in presence of varying concentrations of the universal tyrosine kinase substrate poly-(Glu_4_-Tyr). Representative Coomassie-stained (CS) gels and autoradiographs (A) are shown **C**) Western blot of Sut1 with an anti-phosphotyrosine PY99 antibody. *In vitro* phosphorylation assays were performed with and without ATP and with heat-denatured Sut1. In addition, an assay with bovine serum albumin (BSA) and an assay with the Ser/Thr kinase ArnD were performed as negative controls. Next to the blot image, an image of the corresponding Ponceau-stained membrane is shown. This experiment was performed three times and results were reproducible; a representative result is shown here.

In order to determine the identity of the autophosphorylated residues of Sut1, we tested whether the protein kinase could be recognized by a phosphotyrosine antibody (PY99) (**Figure 4B**). Before performing the western blotting experiments, Sut1 phosphorylation assays were performed in three different conditions *in vitro*: with the addition of ATP, without the addition of ATP or upon adding heat-denatured kinase. In all conditions, including the absence of ATP or upon kinase denaturation, similar signals were obtained, pointing to the prior presence of phosphotyrosines in Sut1. The Ser/Thr kinase ArnD and bovine serum albumin (BSA) were included as negative controls in these experiments; they showed no signals as expected. Overall, these results indicate that Sut1 autophosphorylation occurs on tyrosine residues and that it also occurs upon heterologous expression in *E. coli*.

The above observations were further confirmed by performing mass spectrometry analysis of Sut1, either as purified from *E. coli* or after performing the *in vitro* phosphorylation reaction. In both conditions, Sut1was found to be phosphorylated on two tyrosine residues, namely Tyr89 and Tyr157. None of these tyrosine residues are conserved in other archaeal Sut1 homologs (**Figure 1**), although Tyr157 is located close to the putative catalytic site.

### Site-directed mutagenesis of Sut1

As Sut1 is considered to belong to a novel type of kinases, the catalytic motifs are unknown. In order to identify key residues for kinase activity, Sut1 mutant proteins were prepared and tested *in vitro* for phosphorylation activity (**Figure 5**). Given the previously observed autophosphorylation activity and the general trend that tyrosine kinases are typically activated by autophosphorylation on tyrosine residues [19], Sut1 mutants were constructed in which one or more tyrosine residues were substituted with phenylalanines. This included Tyr157, which is a non-conserved tyrosine found to be phosphorylated in the previously described mass spectrometry analysis, and Tyr161, which is conserved in other archaeal Sut1 homologs and is located adjacent to the putative catalytic Asp160 residue. Mutants were also created for tyrosine residues in the C-terminal tyrosine cluster (Tyr189Phe and Tyr183Phe-Tyr189Phe-Tyr193Phe). All tyrosine mutants displayed diminished kinase activity relative to the wild type, with relative differences (**Figure 5**), suggesting that these tyrosine residues are involved in phosphorylation activity and that some indeed serve as autophosphorylation sites.

**Figure 5.**
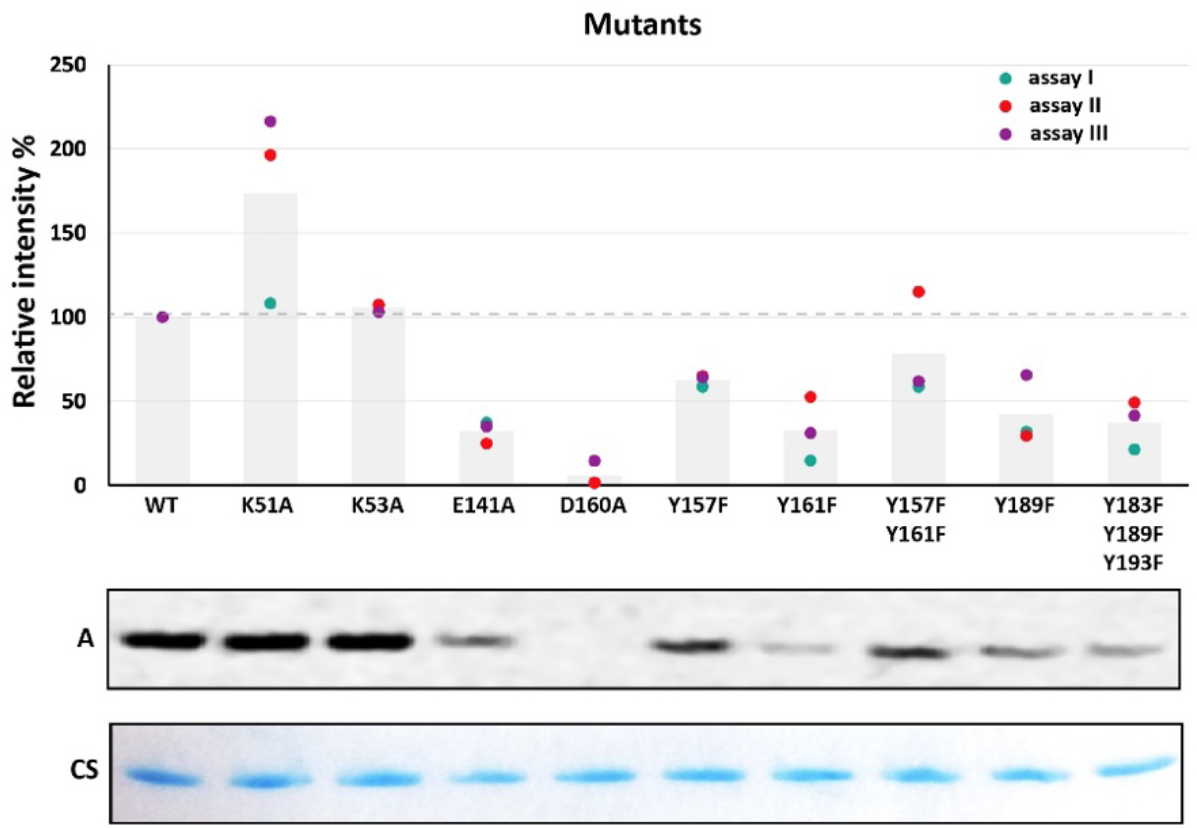
Autophosphorylation activities of different Sut1 mutants. A representative Coomassie-stained (CS) gel and autoradiograph (A) (phosphorimage) are shown. Histograms represent the average of three independent assays, indicated by coloured dots. Relative intensities were estimated using Personal Molecular Imager PMI software (BioRad).

To identify the possible catalytic motifs of Sut1, additional mutants were constructed, including Lys51Ala and Lys53Ala. Both lysine residues are embedded in a Walker A-like motif (**Supplementary Figures 2** and **3**), which is the ATP-binding P-loop in bacterial type tyrosine kinases [8], in which a lysine residue typically plays a key role in ATP anchoring. However, no clear effect on the autophosphorylation activity was observed for these mutants (**Figure 5**), thus revoking the assumption that this is a Walker motif. In contrast, for Asp160Ala or Glu141Ala mutants, a clear negative effect was observed on the phosphorylation activity. While the attenuation was strong for Glu141Ala, the activity was completely lost for the Asp160Ala mutant (**Figure 5**). These observations demonstrate that both residues, which are highly conserved in archaeal Sut1 homologs (**Figure 1**), play a critical role in the catalytic reaction.

### Phosphoproteomic analysis of a *Sut1* disruption strain

It has been previously shown that a gene disruption strain of *Sut1* does not display obvious growth defects [20]. To further unravel the function of Sut1 *in vivo*, and to attempt to identify putative protein targets, phosphoproteomic analyses were performed in the *Sut1* gene disruption strain MW382 and its parental strain MW001 (**Supplementary Tables S1** and **S2**). A total of 1751 proteins were detected in both strains, with 59 and 22 proteins being detected exclusively in MW001 or MW382, respectively, supporting the technical validity of the experiment. This set of identified proteins was filtered based on a two-fold change in their abundance ratio upon comparing both strains and with p < 0.05 cut-off p-value. This filter threshold generated a set of 150 differentially expressed proteins, of which 116 were downregulated in MW382 while 34 were upregulated. The observation that only a small fraction of the detectable proteome is differentially expressed in the Sut1 disruption strain, underscores the previous lack of observing effects on cellular physiology. Differentially expressed proteins belong to various functional categories. Interestingly, several putative transcription factors were found differentially expressed with a DNA-binding protein predicted to encode a ArsR-like transcription factor being one of the most significantly upregulated proteins in the Sut1 disruption strain.

With regards to the phosphorylation events detected in the experiment, 123 unique phosphorylated residues were identified in both strains. The number of detected phosphorylation sites is much lower than previously found in strains in which one or both phosphatases were deleted [3]. Interestingly, among the phosphorylated residues, 11 of these were phosphotyrosines. The abundance ratios of phosphorylation (MW382/MW001)_mod_ were normalized to the abundance ratios of proteins (MW382/MW001)_prot_ in the input (**Supplementary Tables S1** and **S2**).

We assume that due to the dynamic nature of protein phosphorylation, only a small fraction of phosphosites could be detected in the presence of active phosphatases. However, the observation of differential abundances of phosphosites, both tyrosines and serines/threonines, points to Sut1 being involved (in-)directly in different phosphorylation-mediated signal transduction network in *S. acidocaldarius*. The RNA polymerase subunit Saci0693, the thiamine pyrophosphate-containing enzyme Saci0763 and two conserved archaeal proteins Saci1435 and Saci0882 (**Supplementary Table 1**) appear from our data to be potential targets for tyrosine phosphorylation by Sut1, although this needs to be specifically validated by future studies.

### Sut1 phosphorylates the ArsR-family regulator Saci0006 *in vitro*

In the context of identifying putative Sut1 phosphorylation substrates, we also considered a group of 17 transcription regulators that were previously found to be phosphorylated on at least one tyrosine [3]. One of these, a winged-helix-turn-helix (wHTH) DNA-binding protein predicted to be a ArsR-family transcription regulator and encoded by the gene *Saci_0006* was found to be phosphorylated on Tyr67 and Tyr69 *in vivo*. We experimentally assessed whether the Saci0006 protein is a substrate for Sut1 and we found a phosphorylation signal when performing an *in vitro* kinase reaction (**Figure 6**). Finally, to investigate the effect of mutating the C-terminal tyrosine cluster of Sut1 on substrate phosphorylation, Saci0006 was also tested for phosphorylation by the Tyr183Phe-Tyr189Phe-Tyr193Phe triple mutant of Sut1. Besides losing a large part of the autophosphorylation, the cross-phosphorylation activity of the Sut1 triple mutant was also clearly debilitated, emphasizing the role of autophosphorylation in the tyrosine cluster to achieve functional kinase activity (**Figure 6**).

**Figure 6.**
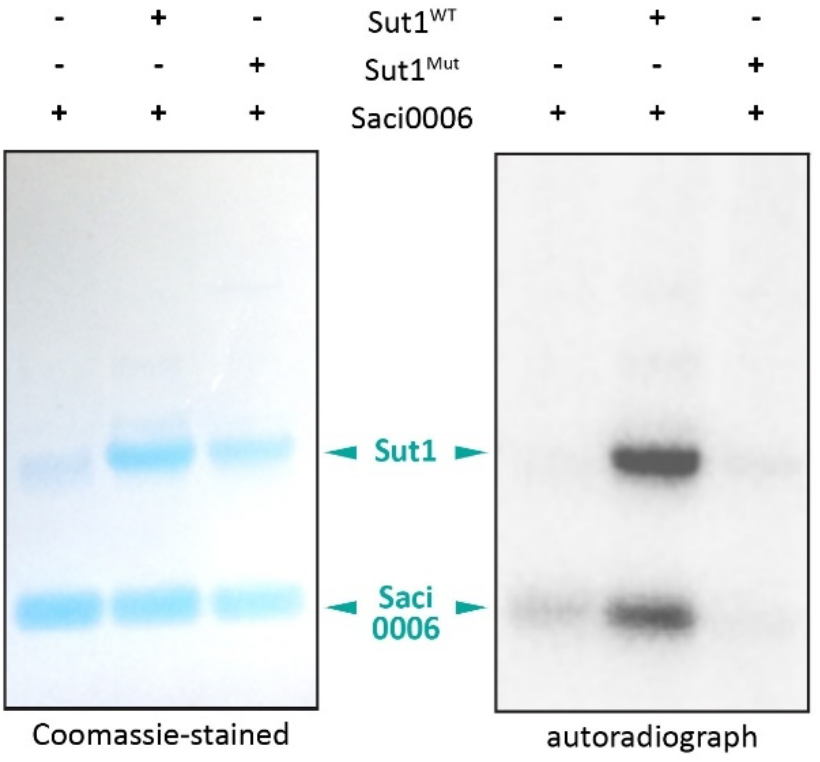
Sut1 phosphorylates Saci0006. Autoradiograph of the phosphorylation assay and corresponding Coomassie-stained (CS) gel of Sut1 WT and mutant (Tyr183Phe-Tyr189Phe-Tyr193Phe) (23.6kDa) with Saci0006 (9.9 kDa).

## Discussion

Tyrosine phosphorylation has been reported to extensively occur in *Sulfolobus spp*. [3]. In this work, we present pioneering steps in identifying a tyrosine kinase in *S. acidocaldarius*, which is to our knowledge the first tyrosine kinase to be reported thus far in archaea. Sut1 represents a novel protein kinase family as its sequence features do not correspond to those of the typical or atypical Hanks-type kinases or to the bacterial tyrosine (BY) kinases. Sut1 is conserved in archaea among various phyla (**Figure 1**), which points towards a possible important role, although the *S. acidocaldarius* homolog does not appear to play a global essential role in the regulation of cellular physiology.

Previously, the *S. islandicus* Sut1 homolog encoded by *SiRe_1057*, did not show any kinase activity [12]. Therefore, a first step in the characterization of this kinase was to obtain it in an active form *in vitro*. Here, we succeeded in purifying an active kinase and upon testing different reaction conditions, the detection of the kinase activity was optimized, resulting in clear phosphorylation signals using [γ-^32^P] ATP. However, the specific activity of the kinase remained relatively low impeding quantification of enzymatic kinetics. This could either be attributed to a small proportion of the protein preparation being active due to folding issues during heterologous expression or due to the requirement of additional cofactors such as a cognate activator (*e*.*g*. transmembrane sensor protein), similarly as found for bacterial tyrosine kinases [8].

The low enzymatic activity was a limitation to obtain consistent mass spectrometry data for the identification of phosphorylation sites. Instead, the use of a universal tyrosine kinase substrate poly-(Glu_4_,Tyr) peptide substrate was a good alternative to test the tyrosine phosphorylation activity of Sut1. Following the approach used in previous studies of other kinases [17][18] phosphorylation signals were clearly detected for poly-(Glu_4_Tyr) by Sut1. The Ser/Thr specific kinase ArnD did not phosphorylate poly(Glu_4_,Tyr), thus confirming the specificity of tyrosine phosphorylation generated by Sut1. Further, when mutating selected tyrosine residues, the autophosphorylation activity of Sut1 was significantly reduced, especially for the tyrosine cluster mutant Tyr183Phe-Tyr189Phe-Tyr193Phe, indicating that these are important autophosphorylation sites. This feature resembles that of the bacterial tyrosine kinase family [8].

We were also able to identify the crucial role of aspartate Asp160 in the “IIDYD” motif, which is highly conserved in Sut1 and its archaeal homologs, for enzymatic catalysis. Mutating this residue leads to a so-called “dead” kinase. Based on the structural homology model that was generated, this motif seems to correspond to the highly conserved “HRD” motif in the Hanks-type protein kinases; hence it can be postulated to be involved in metal cation binding and thus in the phospho-transfer enzymatic reaction. This is in consonance with different protein kinase families, which normally rely on an invariant aspartate for their catalytic activity among different patterns of conserved motifs. Likewise, it is demonstrated that Glu141 in the conserved motif “HK/GEL” is essential for kinase activity. This would correspond to the “DFG” motif in Hanks-type protein kinases. Moreover, the absence of any effect for mutating Lys51 on the autophosporylation activity, indicates that it is not a catalytic lysine in a Walker A like motif. Instead, it is rather a part of the glycine rich loop “GxGxxG” which is highly conserved in Sut1 archaeal homologs and in other protein kinases [4]. As such, we describe a novel tyrosine kinase family and outline its essential putative catalytic motifs, which seem to be different with respect to traditional kinase families. To our knowledge, this is the first tyrosine kinase to be described in archaea. However, given the extent of tyrosine phosphorylation observed in *Sulfolobus* spp., it is unlikely that Sut1 is the only tyrosine kinase that contributes to cellular signaling.

The phosphoproteomic analyses performed provide preliminary indications of the possible functional role of Sut1 in *S. acidocaldarius*. A set of 150 proteins were differentially expressed in the kinase disruption strain. Those differentially expressed proteins are involved in diverse metabolic processes. Some classes are clearly better represented, like transporters (ABC transporters, sugar transporters) and energy metabolism related enzymes (dehydrogenases). One of the proteins that was upregulated in the Sut1 disruption strain was the transcription regulator Saci0006, which we could show that it gets phosphorylated *in vitro* by Sut1. This confirms that Saci0006 is likely an endogenous substrate of the kinase. Our phosphoproteomic data reveals other potential substrates for Sut1.

## Materials and Methods

### Bioinformatic analysis and structural homology modelling

Multiple sequence alignments were performed using Clustal Omega [21] and output figures were generated using the BoxShade server. Sequence logos were generated using the WebLogo server [22]. A structural homology model was retrieved from AlphaFold [9] and the ligand transplants were predicted using Alphafill [11]. Structures were visualized using PyMol (The PyMOL Molecular Graphics System, Version 1.2r3pre, Schrödinger, LLC).

### Microbial strains and growth conditions

*S. acidocaldarius* strains MW001 and MW382 (*Saci_1289*::*pyrEF*) [20] were cultivated in Brock basal medium [23] adjusted to a pH of 3.5 and supplemented with 0.4% (w/v) sucrose and 0.5% (w/v) NZ-amine. For the MW001 uracil-auxotrophic strain, 1 mg/ml of uracil was added to the medium. Cultures were incubated in a shaking incubator at 75°C and growth was assayed by measuring the optical density at 600 nm (OD_600_).

*Escherichia coli* strain DH5a (Gibco) was used for propagation of plasmid DNA constructs and *E. coli* strain Rosetta (DE3) (Novagen) was used for heterologous protein overexpression. *E. coli* strains were cultivated in Lysogeny Broth (LB) medium in a shaking incubator at 37°C. According to the strain and plasmid used, 60 μg/ml kanamycin and/or 25 μg/ml chloramphenicol were added to the medium.

### Cloning, site-directed mutagenesis and protein purification

The *Sut1* gene (encoded by *Saci_1289*) was amplified by PCR from genomic DNA of *S. acidocaldarius* with oligonucleotide primers EP331 and EP332 (**Supplementary Table S3**) and subsequently cloned into the pET28b (Novagen) plasmid vector using NdeI and BamHI restriction sites using classical restriction-ligation cloning, generating pET28b*xSut1*. This construction enables expression of the protein with an N-terminal 6x His tag.

Site-directed mutagenesis was performed for pET28b*xSut1* by full plasmid amplification with two mutagenic primers as described by [24]. All mutant constructs were transformed into *E. coli* Rosetta (DE3) for heterologous protein production. An overview of all oligonucleotides used in this study is presented in **Supplementary Table S3**.

For the wild-type and mutant Sut1 proteins, heterologous protein expression was performed by incubating the culture at 37°C until an OD_600_ of 0.7 was reached, followed by adding 0.4 mM isopropyl ß-galactopyranoside (IPTG) to induce expression. Subsequently, the culture was further cultivated for either 4 hours at 37°C or overnight at 16°C. Cells were harvested by centrifugation at 13689 *g*, washed in 0.9% NaCl and resuspended after centrifugation in lysis buffer (20 mM phosphate buffer [pH 7.4], 500 mM NaCl, 40 mM imidazole, 1 mM Pefablock protease inhibitor). Lysis was performed by sonication for 10 minutes at 25% amplitude at 4°C (Vibra Cell sonicator). Afterwards, cell debris was removed by centrifugation at 5000 *g*. The protein was then purified from the supernatant by immobilized metal affinity chromatography (IMAC) as described in [25]. Samples of the elution fractions were analyzed by SDS-PAGE on a 4-12% Bis-Tris gel (Invitrogen) to check the presence and purity of the protein. Fractions with pure protein were pooled and dialyzed against phosphate buffered saline (PBS) buffer [pH 7.4]. Finally, the obtained protein concentration was measured by absorption at 280 nm using a NanoDrop spectrophotometer (Thermo Scientific).

### In vitro phosphorylation assay

Kinase activity assays were performed using 5.5 µM of pure protein in the following reaction buffer: 25 mM Tris-HCl [pH 7.4], 100 mM NaCl, 10 mM MgCl_2_, 1 mM Na_3_VO_4_, 1 mM DTT, 100 µM ATP and 1.5 µCi [γ-^32^P]ATP (3000 Ci mmol^−1^, PerkinElmer), unless otherwise stated. Assays to test different cations were performed in a different reaction buffer (20 mM MES, 50 mM NaCl, 2 mM DTT, 100 µM ATP and 1.5 µCi [γ-^32^P] ATP) to avoid precipitation of metals. In some assays, poly-(Glu_4_,Tyr) sodium salt (Glu:Tyr (4:1)), molecular weight 5,000-20,000(Sigma-Aldrich) was added as a substrate in a range between 2 µg and 6 µg for the total reaction mixture, while in other assays 2.6 µM of heterologously purified Saci0006 protein was added as a substrate.

Reaction mixtures were prepared on ice, followed by an incubation at 70°C during 1 hour, or as otherwise stated. To stop the reactions, NuPAGE LDS loading dye (ThermoFisher) was added. Samples were then incubated at 95°C during 5 minutes and separated by SDS-PAGE using 4-12% Bis-Tris gels (Invitrogen), stained for 1 hour using BioSafe-Coomassie stain (BioRad), and finally de-stained in distilled water for 2 hours. Afterwards, the gel was dried in a heat gel dryer (BioRad) for 40 minutes. Subsequently, the gel was exposed overnight to Storage Phoshor Screen (Kodak), after which images were obtained by scanning with a Personal Molecular Imager (BioRad). The densitometric quantification of the signals was performed using Fuji [26]. Calculations of the specific activity were made following a method adapted from [27].

### Western blotting

Kinase assays were performed as mentioned above. After separation by SDS-PAGE, proteins were blotted on a 0.45 µm nitrocellulose membrane (Bio-Rad), followed by verifying the transfer by Ponceau staining. Next, the membrane was blocked using 1% bovine serum albumin for 1 hour and washed with Tris-buffered saline (TBS) buffer (50 mM Tris-Cl [pH 7.5], 150 mM NaCl) twice for 5 minutes. Afterwards, the membrane was incubated with a 1:500 dilution of the primary phosphotyrosine antibody PY99 (Santa Cruz) overnight at 4°C. After washing with TBS buffer containing 0.1% Tween 20 (TBST) three times for 10 minutes each, the membrane was incubated with a 1:10000 dilution of the IRDye 680RD goat anti-mouse IgG secondary antibody (LI-COR) during 1 hour at room temperature. Membranes were washed again using TBST thrice for 10 minutes. The resulting signals were detected by scanning the membrane using an Odyssey CLx imaging system (LI-COR) and its *ad hoc* software.

### Mass spectrometry and phosphoproteomics

For (phospho-)proteomic analysis, *S. acidocaldarius* strains MW001 and MW382 were grown in 500 ml cultures as mentioned before. At OD_600_ 0.8, cells were harvested by centrifugation during 20 minutes at 13000 x *g* (Sorvall, SLA-3000 rotor). Cell pellets were then washed with 10 ml cold 0.9% NaCl solution and resuspended in 5 ml lysis buffer (PBS [pH 7.6], 5 mM NaF, 5 mM sodium orthovanadate, 5 mM Pefablock and 1 µg/ml leupeptine). Cells lysis was done by sonicating at 20% amplitude for 15 min followed by centrifugation at 10000 x *g* for 40 min at 4°C. The supernatant was recovered and then centrifuged at 10000 x g for 20 min at 4°C. Samples were aliquoted, flash-frozen in liquid nitrogen and stored at -80°C until further processing. The protein content was estimated using the Bradford method, according to manufacturer’s instructions.

A total of 3.2 mg protein, divided into 400-µg aliquots, was precipitated by adding 15% (v/v) trichloroacetic acid TCA and incubated on ice for 30 minutes. Next, samples were centrifuged for 15 minutes at 20000 x *g* and 4°C, followed by discarding the supernatants and resuspending each pellet in 50 mM ammonium bicarbonate [pH 8.5]. Overnight digestion was performed by adding trypsin (ratio 1:50, Promega) at 37°C while shaking. Another 3 hours incubation with fresh trypsin was conducted the day after to ensure a complete digestion of samples. After centrifuging for 10 minutes at 10000 x *g*, all peptide-containing supernatants from the same extract were pooled and dried in a SpeedVac. Finally, pellets were resuspended in 3.5% acetonitrile, 0.1% TFA and peptide content was estimated with a Pierce™ Quantitative Colorimetric Peptide Assay (Thermo Scientific) according to the manufacturer’s instructions. A total of 2.5 mg of the generated peptides were loaded on top of TiO_2_ beads (GL Sciences) and processed according to the manufacturer’s instructions, except for the elution step, which was performed by two sequential steps with 50 µl of fresh 5% (v/v) ammonia. The resulting samples were then dried in a SpeedVac and pellets were resuspended in 3.5% acetonitrile (ACN), 0.1% trifluoroacetic acid (TFA).

A volume of 6 µl phosphopeptides or 500 ng of peptides were directly loaded onto reversed-phase pre-column (Acclaim PepMap 100, Thermo Scientific) and eluted in backflush mode. Peptide separation was performed using a reversed-phase analytical column (Acclaim PepMap RSLC, 0.075 × 250 mm (Thermo Fisher Scientific)) with a linear gradient of 4-27.5% solvent B (0.1% FA in 98% ACN) for 100 minutes, 27.5-40% solvent B for 10 minutes, 40-95% solvent B for 1 minute and holding at 95% for the last 10 minutes at a constant flow rate of 300 nl/min on an EASY-nLC 1000 RSLC system. The peptides were analyzed by an Orbitrap Fusion Lumos tribrid mass spectrometer. The peptides were subjected to NSI source followed by tandem mass spectrometry (MS/MS) in Fusion Lumos coupled online to the nano-LC. For total proteome comparison, intact peptides were detected in the Orbitrap at a resolution of 120,000. Peptides were selected for MS/MS using HCD setting at 30; ion fragments were detected in the Ion Trap. A data-dependent procedure that alternated between one MS scan followed by MS/MS scans was applied for 3 seconds for ions above a threshold ion count of 5.0E3 in the MS survey scan with 60.0s dynamic exclusion. The electrospray voltage applied was 2.1 kV. MS1 spectra were obtained with an AGC target of 4E5 ions and a maximum injection time of 50ms, and MS2 spectra were acquired with an AGC target of 3E4 ions and a maximum injection time of 35ms. For MS scans, the m/z scan range was 350 to 1500. For phosphopeptides, the MS2 spectra were acquired in the Orbitrap at a resolution of 60,000 with an AGC target of 1.0E5 ions and a dynamic injection time to maximize sensitivity.

The resulting MS/MS data was processed using Sequest HT search engine within Proteome Discoverer 2.4 SP1 against *S. acidocaldarius* protein database obtained from Uniprot (2221 entries). Trypsin was specified as cleavage enzyme allowing up to 2 missed cleavages, 4 modifications per peptide and up to 5 charges. Mass error was set to 10 ppm for precursor ions and 0.1 Da for fragments ions. Oxidation on Met, phosphorylation on Ser, Thr and Tyr were considered as variable modifications. The false discovery rate (FDR) was assessed using Percolator and thresholds for protein, peptide and modification site were specified at 1%. For total proteome comparison, abundance ratios were calculated by Label Free Quantification (LFQ) of the precursor intensities within Proteome Discoverer 2.4 SP1. For phosphopeptides, ratios were calculated as the median of all possible pairwise peptide ratios calculated between replicates using only phospho containing peptides. The application then uses the paired (background-based) method to calculate the p-value.

## Acknowledgements

This article is dedicated to the memory of Pierre Cornelis. We are grateful to Ann-Christin Lindås for the gift of heterologously purified Saci0006 protein and to Sonja-Verena Albers for the gift of *S. acidocaldarius* strains MW001 and MW382. We thank Karl Jonckheere for technical assistance with protein purification work. This study was supported by the Vrije Universiteit Brussel (VUB) Research Council (Strategic Research Program SRP91) and the Research Foundation Flanders (FWO-Vlaanderen) (EP and GJG) (Research Projects G021118N and G062820N to EP).

## Author contributions statement

HRM and EP conceived the work; HRM and GJG performed the experiments and collected data; IB conducted the mutagenesis experiments, SPdR and DV executed and analyzed the proteomic experiments. HRM, GJG, BS, WV and EP analyzed data and HRM and EP wrote the manuscript. All authors approved the final article.

