## Supplementary for "Protein kinase Sut1 from the Crenarchaeon *Sulfolobus acidocaldarius* displays tyrosine phosphorylation activity"

### Supplementary material

```
                                                                      ↓
Sut1            -ENFILMEYIRGRHL-TRQENIE------------T-----IRKILMKARMLELKGIEHKELSRPYKNVLVNEKDA----
Saci1193        -PPAIVMEYMEGGDL-RQIITNNEYSTLRHSSRWPEIVSIMFSKIADAIIHVNKNGYVHCDIK--PSNILLNGKLPK---
Saci1041        -PPAIIMEFMDGGTI-ADLINE-----IRSSPYFQYIVKAIIREVGLALKYLHKRGYVHLDVK--PRNIFLSVIPP----
Saci1694        -PPYVAMEYMGNGTV-KDLLLRDE---YFYSVRWKKLVYRIIFQVARGLLYLHSEGYVHLDVK--PSNIFIESVKG----
Saci0965        -KNILIMEFIGDNGIRAPLLKEL-----RSEDVNEDLYKTIID---QVVIMANKAELVHGDLS--EYNIMVFDGKP----
Saci0850        IQFLIVMEYIEGTTL-RDIFNQG------------RNLRELGKEMGLMISLLHKANIIHGDLT--TTNMIYTDEGL----
Saci1181        -PPAIIMEFMEGGTL-FDLISRVD---LVQSKYWQYIVKLVIKEIAKALTFLHKRGYVHLDVK--PQNIFLKEKIN----
Saci1664        -KRVLIMEYVKGYKV-TSEEAKR----IMRAPDLSY---MVFRTF---MVMLLTKEYFHADPH--PGNIALDEKGNLILY
Saci0796        -LNAIAMEYIYGKEL-YKIEISDP---E---SALNDI----LSTVRI---AYKECNIIHGDLS--EYNILYSLDS-----
Saci1869        -PPAIVMEYMEGGTL-ADLMRNDI---VVNSSHWGDIVKVVARKVAKALDYIHSSGYVHLDVK--PQNIFFSAPVG----
                    : **::  .  :                                          * :      *:
                                                                      Subdomain VIb
```

**Supplementary Figure 1**. Partial sequence alignment of protein kinases in *S. acidocaldarius.* Protein sequence alignment of ten protein kinases in *S. acidocaldarius,* with indication of the conserved catalytic subdomain VIb. The catalytic aspartate is indicated with a red arrow. Identical residues and similar residues are black- and grey-shaded, respectively.

```
Sut1      MDADMKSIDIKYFIYPLYSAEIEKELRANGINSLYSFGYVKLRPDINVIGKGKTGIVALF  60
CpsD      ------------MPTLEIAQKK---L---EFIKKAEEYYNALCTNIQLSG----------  32
PtkA      MA--LRKTRGSWMRRTVIAMT--------EPKSLNSEQYRTIRTNIEFASVD--------  42
CapB      MA--KKK----STISPLYVHD--------KPKSTISEKFRGIRSNIMFSNAE--------  38
                                          .  .  :  :  :* . .

                         Walker A              Walker A´
Sut1      DENKVIKIRRTD------------------SPKETLKIEAKIQQLAYPVSPKVYLYGENF  102
CpsD      DKLKVISVTSVNPGEGKTTTSVNIARSFARAGYKTLLIDGDTRN---SVMSGFF------  83
PtkA      RQMKTVMITSACPGEGKSTTAANLAVVFAQQGKKVLLIDADLRK---PTVHTAF------  93
CapB      NEIKSLLITSEKSASGKSILSANIAVTYAQAGYKTLIIDGDMRK---PTQHYIF------  89
           : * : :                         :.* *:.. ::    .    :

Sut1      ILMEYIRGRHLTRQENIETIRKILMKARMLELKGIEHKELSRPYKNVLVNEKDAYIIDYD  162
CpsD      -----------KSREKITGLTEFLSGT--------------ADLSHGLCDTNIENLFVVQ  118
PtkA      -----------HL-ENMIGLTTVLLKK--------------SSLEQAVQASNEKYLDVLT  127
CapB      -----------DL-PNNSGLSNLIINK--------------TTYSDSIKETRVENLNVLT  123
                         :   :  .:                     .. :   .   :

                                         Walker B
Sut1      SATIKPNPKNVTSILSWFKRYEL--AKMYSLGY----PL-------------EKIILLL-  202
CpsD      SGTVSPNPTALLQS-KNFNDMIETLRKYFDYIIVDTAPIGIVIDAAIITQKCDASILVTA  177
PtkA      SGPIPPNPAELLSS-KWMKELADEACAAYDMVIFDTPPILAVADAQILGNVADGSVLVIS  186
CapB      AGPTPPNPSELIAS-SKFATIFNELLNHYDFIVIDTPPINTVTDAQVYARIVKNCVLVID  182
          :.   ***  :    . :          :.       *:             .  :*:

                                        <Tyrosine rich tail>
Sut1      ------------------------------------------------------        202
CpsD      TGEVNKRDVQKAKQQLEQTGKLFLGVVFNKLDISVDKYGVYGFYGNYGKK----        227
PtkA      SGKTEKEQAAKAKEALESCKSKLLGAIMNGKKLS--KHSEYGYYGNKENFIQNK        238
CapB      AEKNNKSEVKKAKGLLTKAGGKVLGAVLNKMPID--KNSSYYYYYGED------        228
```

**Supplementary Figure S2.** Sequence alignment of Sut1 and bacterial tyrosine (BY) kinases. Protein sequence alignment of Sut1 from *S. acidocaldarius* with the BY-family kinases CpsD from *Streptococcus pneumoniae* (Nourikyan et al., 2015), PtkA from *Mycobacterium tuberculosis* (Bach et al., 2009) and CapB from *Staphylococcus aureus* (Olivares-Illana *et al*.,2008). Key conserved motifs of the BY family Walker A, Walker A' and Walker B are indicated by red rectangles. Multiple tyrosine residues in Sut1 in the C-terminal domain are indicated in blue.

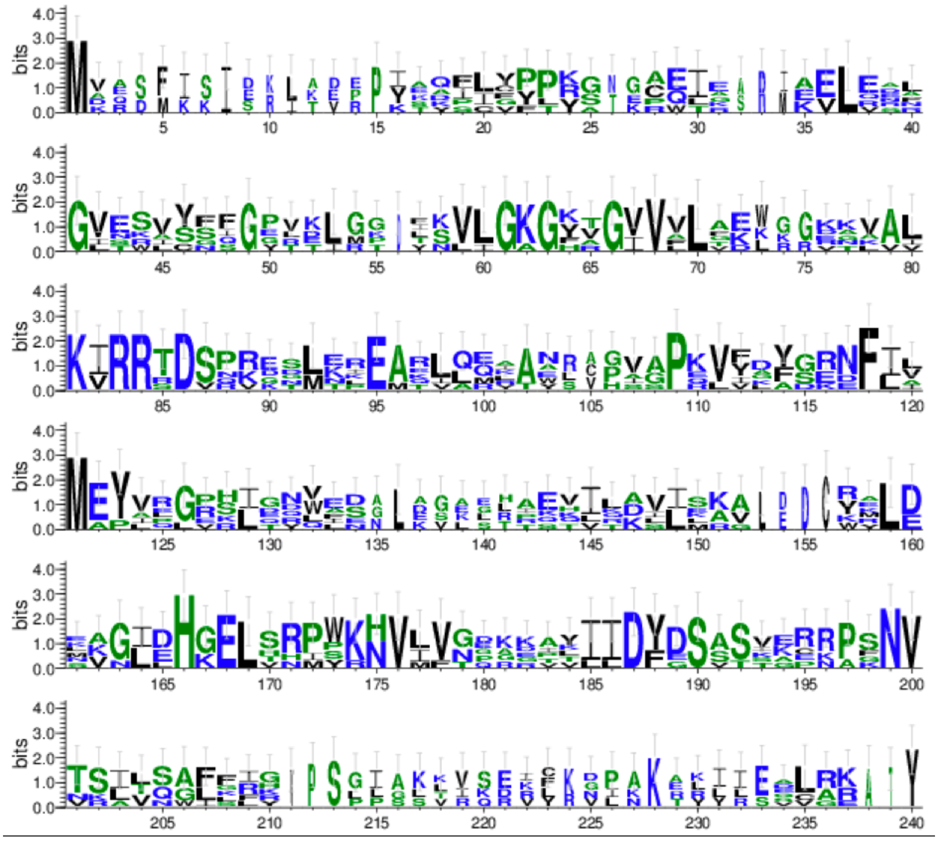


**Supplementary Figure S3**. Sequence conservation in archaeal Sut1 homologs. This sequence logo was generated for the alignment of putative Sut1 homologs that are presented in **Figure 1**. The x-axis indicates the position, and the y axis represents the probability of occurrence of an amino acid. The sequence logo was generated using WebLogo (Crooks *et al.*, 2004).

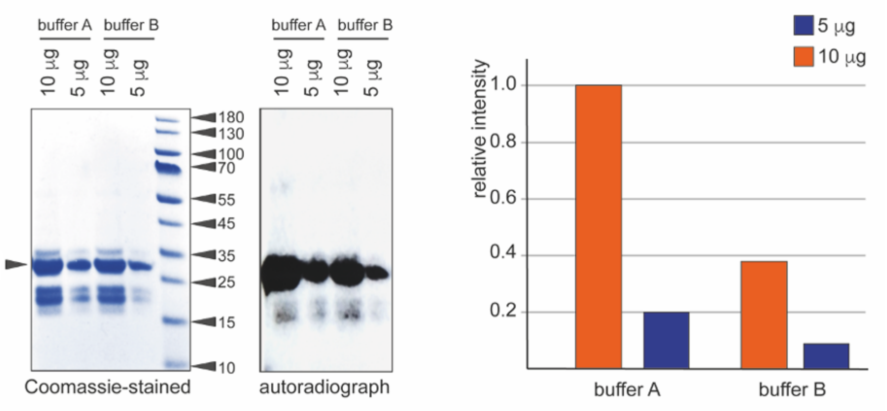


**Supplementary Figure S4.** Effect of reaction buffer composition on Sut1 autophosphorylation activity *in vitro*. Qualitative and quantitative results of an *in vitro* phosphorylation assay of Sut1 (23.6 kDa). The Coomassie-stained gel and corresponding autoradiograph are shown side-by-side, with indication of the molecular weights of the proteins ladder in kDa. Relative intensities were calculated using a Personal Molecular Imager PMI software (BioRad). Besides full-length Sut1 additional protein species are observed, which are possibly degradation products of the protein.

**Supplementary Table S1.** List of proteins with a detected phosphotyrosine in the phosphoproteomic analysis, with a lower level of abundance in the *Sut1* disruption strain (MW382) with respect to the parental WT strain (MW001). Proteins with a significant abundance ratio are shaded in orange.

| **Protein** | **Description** | **Peptides with phosphosite (indicated in red)** | **Abundance ratio (MW382/MW001)** |
|---|---|---|---|
| Saci_0693 | DNA-directed RNA polymerase subunit B | R**Y**$^{1076}$VCPVHGDKS | 0.072 |
| Saci_0763 | Thiamine pyrophosphate-requiring enzyme | K**Y**$^{50}$EIKPNLCNSTRE | 0.012 |
| Saci_1435 | Conserved archaeal protein | CEKK**Y**$^{45}$ NAF | 0.3 |
| Saci_0882 | Conserved archaeal protein | R**Y**$^{15}$EIIHDILAQCENGAKK | 0.43 |
| Saci_1249 | Gylcosyl transferase group 1 protein | YKEGIYNESHGIEK**Y**$^{106}$ | 0.782 |
| Saci_1401 | Thermosome subunit alpha | KDATPEDLG**Y**$^{352}$AELVEERR | 0.804 |

**Supplementary Table S2.** List of proteins with a detected phosphotyrosine in the phosphoproteomic analysis, with a higher level of abundance in the *Sut1* disruption strain (MW382) with respect to the parental WT strain (MW001). Proteins with a significant abundance ratio are shaded in orange.

| **Protein** | **Description** | **Peptides with phosphosite (indicated in red)** | **Abundance ratio (MW382/MW001)** |
|---|---|---|---|
| Saci_0745 | Conserved archaeal protein | SFLDEEAR**Y**$^{352}$ | 5.6 |
| Saci_1690 | Alcohol dehydrogenase | KVNLSQL**Y**$^{270}$SKH | 4.4 |
| Saci_1470 | Quinolinate phosphoribosyl transferase | KMTDI**Y**$^{20}$ FDRT | 3.07 |
| Saci_1116 | Acetyl esterase | RSMVEY**Y**$^{192}$DGYFLTRE | 1.535 |
| Saci_0663 | Histidine--tRNA ligase | R**Y**$^{110}$DEPQFGRY | 1.519 |

**Supplementary Table S3.** An overview of the oligonucleotides used in this study.

| Name | Sequence 5'-> 3' | Purpose |
|---|---|---|
| EP331 | GGAATTCCATATGGATGCAGACATG | Cloning of *Saci_1289* in pET28b |
| EP332 | CGGGATCCCTATAACAATAGAATGATTTTTTC | Cloning of *Saci_1289* in pET28b |
| HM200 | GAAAAAGATGCGTATATAATTGCGTATGATTCA GCAACTATTAAACC | Site-directed mutagenesis of Saci1289 D160A |
| HM201 | GAAAAAGATGCGTATATAATTGCGTATGATTCA GCAACTATTAAACC | Site-directed mutagenesis of Saci1289 D160A |
| HM202 | GTTAAGACCAGATATAAATGTGATAGGAGCTGG AAAAACTGGAATAGTAGC | Site-directed mutagenesis of Saci1289 K51A |
| HM203 | GCTACTATTCCAGTTTTTCCAGCTCCTATCACA TTTATATCTGGTCTTAAC | Site-directed mutagenesis of Saci1289 K51A |
| HM204 | GTTAGAGCTAAAGGGAATAGAACATAAAGCTTT ATCGAGACCCTACAAGAATG | Site-directed mutagenesis of Saci1289 E141A |
| HM205 | CATTCTTGTAGGGTCTCGATAAAGCTTTATGTT CTATTCCCTTTAGCTCTAAC | Site-directed mutagenesis of Saci1289 E141A |
| HM206 | AGATATGAACTTGCAAAAATGTTCTCACTTGGT TATCCTTTAGAAAAAATCA | Site-directed mutagenesis of Saci1289 Y189F |
| HM207 | TGATTTTTTCTAAAGGATAACCAAGTGAGAACA TTTTTGCAAGTTCATATCT | Site-directed mutagenesis of Saci1289 Y189F |
| IB0512 | GACCAGATATAAATGTGATAGGAAAGGGAGCGA CTGGAATAGTAGCTTTATTCGACGAG | Site-directed mutagenesis of Saci1289 K53A |
| IB0513 | CTCGTCGAATAAAGCTACTATTCCAGTCGCTCC CTTTCCTATCACATTTATATCTGGTC | Site-directed mutagenesis of Saci1289 K53A |
| IB0514 | CCATCTTATCTTGGTTCAAAAGATTTGAACTTG CAAAAATGTTTTCACTTGGTTTTCCTTTAGAAA AAATCATTCTATTG | Site-directed mutagenesis of Saci1289 Y183F-Y189F-Y193F |
| IB0515 | CAATAGAATGATTTTTTCTAAAGGAAAACCAAG TGAAAACATTTTTGCAAGTTCAAATCTTTTGAA CCAAGATAAGATGG | Site-directed mutagenesis of Saci1289 Y183F-Y189F-Y193F |
| IB0516 | GTTAATGAAAAAGATGCGTTTATAATTGATTTT GATTCAGCAACTATTAAACC | Site-directed mutagenesis of Saci1289 Y157F-Y161F |
| IB0517 | GGTTTAATAGTTGCTGAATCAAAATCAATTATA AACGCATCTTTTTCATTAAC | Site-directed mutagenesis of Saci1289 Y157F-Y161F |
| IB0518 | GTATTAGTTAATGAAAAAGATGCGTTTATAATT GATTATGATTCAG | Site-directed mutagenesis of Saci1289 Y157F |
| IB0519 | CTGAATCATAATCAATTATAAACGCATCTTTTT CATTAACTAATAC | Site-directed mutagenesis of Saci1289 Y157F |
| IB0520 | GTTAATGAAAAAGATGCGTATATAATTGATTTT GATTCAGCAACTATTAAACCAAATCC | Site-directed mutagenesis of Saci1289 Y161F |
| IB0521 | GGATTTGGTTTAATAGTTGCTGAATCAAAATCA ATTATATACGCATCTTTTTCATTAAC | Site-directed mutagenesis of Saci1289 Y161F |
